# Adult emergence order in a community of cavity-nesting bees and wasps, and their parasites

**DOI:** 10.1101/556456

**Authors:** J Scott MacIvor

## Abstract

Evaluating resource use and overlap through time and space among and within species having similar habitat requirements informs community-level conservation and coexistence, efforts to monitor species at-risk and biological invasions. Many species share common nesting requirements; one example are cavity-nest bees and wasps, which provision nests in dark and dry holes in wood, plant stems, or other plant-based materials that can be bundled together into ‘trap nests’. In this study, the adult emergence order of 47 species of solitary cavity-nesting bees and wasps, and their parasites (total N>8000 brood cells) were obtained from two hundred identical trap nests set up each year (over three years) to survey these populations across Toronto, Canada and the surrounding region. All brood cells collected were reared in a growth chamber under constant warming temperature and humidity to determine species identity, and adult emergence order. This order ranged from 0 to 38 days, with all mason bees (*Osmia* spp.) emerging within the first two days, and the invasive resin bee species, *Megachile sculpturalis* Smith significantly later than all others. Late emerging species i) exhibited significantly greater intraspecific variation in mean emergence day and ii) were significantly larger in body size, compared to early emerging species. Detailing natural history information at the species- and community-level, such as the adult emergence order of coexisting cavity-nesting bees and wasps and their parasites, can inform the timing of deployment of trap nests to support and monitor target species, and refine experimental design to study these easily-surveyed and essential insect communities.

## Introduction

Interspecific partitioning in the timing of similar, critical life-history stages is an evolutionary adaptation used to minimize resource overlap and competition among species (Richards 1927; Schoener 1974; Albrecht and Gotelli 2001; Martin et al. 2004; Taylor et al. 2014). For example, many nest provisioning solitary bee and wasp species share similar nesting location preferences, yet have evolved to partition these resources in order to coexist in time (Wcislo and Cane 1996; Hoehn et al. 2008) and space (Willmer and Corbet 1981; Tylianakis et al. 2005). Intraspecific variation at these critical life-history stages can also be important in determining fitness among individuals within the same species (Bolnick et al. 2011). Variation in emergence from nests as adults may be linked to evolutionary adaptations to local environmental conditions that optimize reproductive and foraging success. In temperate regions, solitary bees and wasps are active for short overlapping periods in a season, which are linked to the availability of preferred resources (Lindsey 1958; Minckley et al. 1994; Leong and Thorp 1999). The remainder of the year is spent as an immature in a nest constructed by the mother (with some exceptions).

Considerable information is available from field and laboratory studies on the lifecycle, incubation period, and adult emergence order of specific solitary bee species managed for pollination in agriculture, such as *Osmia lignaria* Say (Bosch et al. 2000), *Osmia cornifrons* (Radoszkowski) (White et al. 2009), or *Megachile rotundata* (Fabricius) (Tepedino and Parker 1986). However, there are relatively little data on community-wide adult emergence order or knowledge of intraspecific variation in emergence order of most coexisting solitary bees, and especially wasps, many of which are important predators in biological communities (Budrienė et al. 2004; Forrest and Thomson 2011; Fründ et al. 2013). Many solitary bee and wasp species provision nests in cavities above ground (e.g. pithy or hollowed out plant stems, beetle-bored holes in wood), hereafter ‘cavity-nesting bees and wasps’ (Stephen and Osgood 1965; Bohart 1972; Williams et al. 2010). For many cavity-nesting species, artificial nests that represent the natural nesting conditions can be made by gathering ‘nesting tubes’ (e.g. drilled holes in wood, plant stems, or paper cardboard tubes) together, commonly referred to as ‘trap nests’ (Figure 1) (Krombein 1967). Trap nests are widely used to survey cavity-nesting bees and wasps, and their parasites, which include cleptoparasitic bees and wasps, as well as parasitoid wasps, flies, and beetles (Tscharntke et al. 1998; Praz et al. 2008; MacIvor 2017). Identifying nests occurring naturally in the landscape is time consuming and difficult to retrieve in sufficient numbers needed for experiments. On the other hand, rearing cavity-nesting bees and wasps, and their parasites to adulthood from individuals obtained from nesting tubes in trap nests is a useful way to identify species-level interactions and community-level patterns at a local or landscape scale (Staab et al. 2018). Many other studies utilize trap nests to evaluate relationships between species diversity and resource utilization, and in response to environmental change (Yocum et al. 2005; Sheffield et al. 2008; O’Neill et al. 2011; Fliszkiewicz et al. 2012; Fründ et al. 2013). There are applications as well; for example, to support species of concern, the addition of foraging plants (Sheffield et al. 2008) or accelerate the release of bees reared on mass to synchronize with target crops (Bosch et al. 2000).

**Figure 1.**
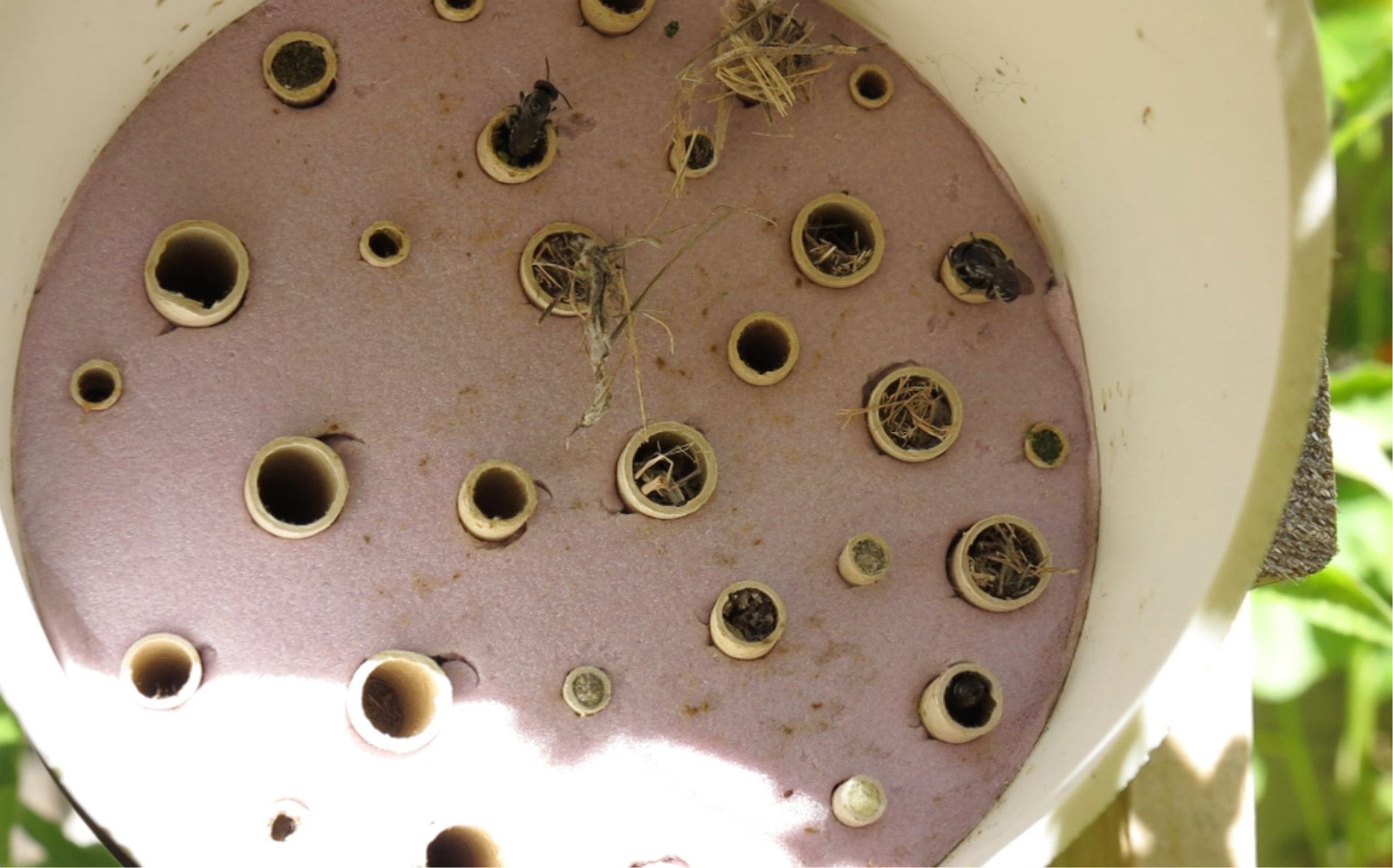
A trap nest used to study cavity-nesting bees and wasps. The cardboard paper ‘nesting tubes’ are inserted into a piece of pink insulation board fitted into a white PVC pipe for support and protection from rain.

In this study, the adult emergence order among and within 47 coexisting species of cavity-nesting bees and wasps, and their parasites were quantified from individuals obtained from trap nests and reared in a laboratory setting at constant temperature and humidity, to represent a spring warming period. The main objective was to map emergence order in this community of cavity-nesting bees and wasps, and their parasites to improve management of these important taxa, for example, to 1) support an abundance of target species, 2) aid in the detection of invasive species, or 3) monitor host-parasite interactions as environmental indicators.

Using these data, I also evaluate two hypotheses. First, that intraspecific variation in emergence time will increase in late emerging species because they incubate longer than early emerging species, and fluctuations in the environment can affect development time (O’Neill et al. 2011). Second, although Bosch and Kemp (2002) noted no relationship between intraspecific body size and adult emergence in the European Orchard bee, *Osmia cornuta*, I predicted body size differences between species in the bee and wasp community would affect interspecific adult emergence order, with larger bodied species emerging significantly later than smaller bodied species, because body size is generally correlated with development time to adult (Garcia-Barros, 2000).

## Methods

The cavity-nesting bees and wasps, and their parasites examined here were obtained from a survey of 200 trap nests set up each year (one per site) from May to October for three years (2011-2013) throughout the city of Toronto and surrounding region. The sites were a minimum 250m apart and spread over an area encompassing approximately 745km^2^. Trap nests were made of white PVC piping that was 10cm in diameter and 28cm in length, a circular faceplate made of insulation board into which 30 cardboard nesting tubes (Custom Paper Tubes, Cleveland, OH) were inserted (10 of each of three tube diameters; 3.4mm, 5.5mm, 7.6mm; all 15cm in length) at one end, and the opposite end was blocked with a PVC pipe cap (Figure 1). Each year, in October, trap nests were collected, each cardboard nesting tube opened, and every brood cell removed. The contents of each brood cell were labeled with a unique identifier and placed into individual cells within 24-cell assay trays with the lid on. A complete incubation process includes a sufficiently long cooling period (Bosch and Kemp 2004), and so all specimens spent the cold season (October – March) in a walk-in fridge kept at a constant 4°C (as in Bosch et al. 2000).

At the end of the cold season, in early April the following year, the assay trays containing all brood cells from the previous year were moved from the walk-in fridge to a sealed growth chamber where temperature and humidity were held constant at 26°C and 60% (Johansen and Eves 1973; Tepedino and Parker 1986). All species overwintered as prepupae (mature post-defecated larvae) except for mason bees (*Osmia* spp.), and a parasite wasp of *Osmia* (*Sapyga centrata* Say) which overwinter as adults. The growth chamber was windowless and kept dark for the duration of the study except when lights were turned on during daily inspection of brood cells to measure adult emergence order (as in Sheffield et al. 2008). Approximately 3% of all cells in the growth chamber were lost to parasitic wasps (*Melittobia chalybii* Ashmead and *Monodontomerus obscurus* Westwood) wasps that emerged early and attacked other larvae still undergoing development. To reduce their depredations, four traps, each consisting of a black light and a bowl filled with water and dish soap, were set up to attract and reduce the number of these minute wasps that emerged and escaped the assay tray (Eves et al. 1980). For each individual bee, wasp, or parasite, the time to emergence was recorded as the number of days from the start of warming (e.g. onset of the incubation period) to the time of development to adulthood and emergence from the cocoon (Owen and McCorquodale 1994; Sheffield et al. 2008). After the three-year survey, a total of 84 species of bee, wasp, and parasite were identified (MacIvor and Packer 2015), but emergence day was recorded for only those species with 5 or more successfully emerged adults with timing accurately recorded, and so 47 species and 8,006 individuals were used for this study (Figure 2). All bees, wasps, and parasites were identified using the collections from the Packer Lab at York University and the Marshall Lab at University of Guelph. All specimens are kept in the MacIvor Lab at University of Toronto Scarborough.

**Figure 2.**
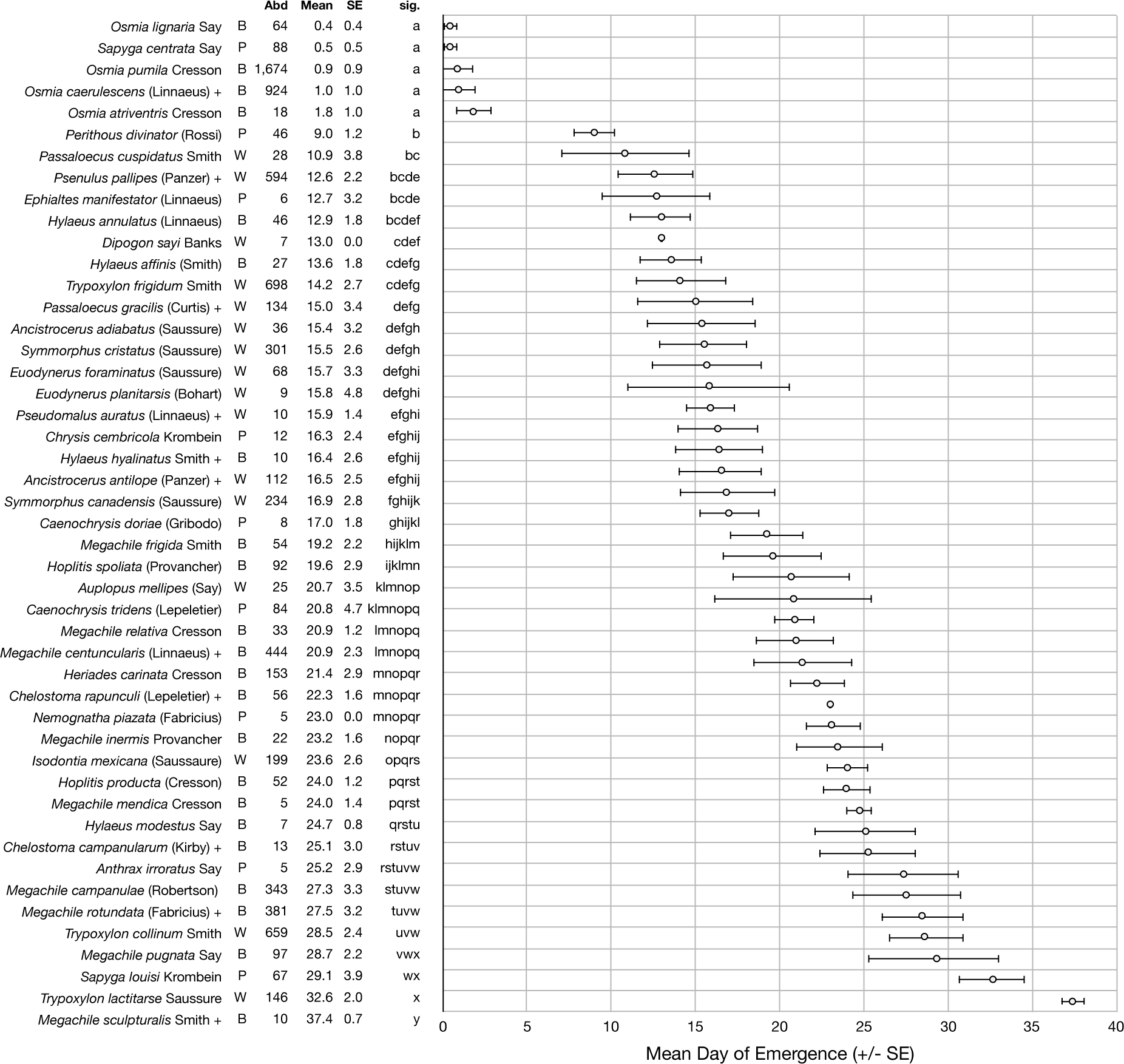
Variation in the mean emergence time for species of bee, wasp, and parasite recorded from individuals taken from trap nests. ‘Taxa’ denotes cavity-nesting bees (B), cavity-nesting wasps (W), and parasites (P). ‘Abd’ is the total number of individuals incubated and emerged successfully. ‘Mean’ is the average number of days taken to emerge and ‘SE’ is the standard error of the mean. Significant differences between species (‘sig’) were given alphabetically where species sharing a letter were not significantly different from one another (α = 0.05). Those species denoted with a “+” are considered introduced to the region.

An analysis of variance (α=0.05) was used to determine if there was a significant difference in interspecific adult emergence order (in days post onset of incubation) among the cavity-nesting bees and wasps, and their parasites. Given my objectives, all males and females per species were grouped together; even though males emerge on average slightly earlier than females, both sexes emerge near the same time to ensure reproduction. A tukey-post hoc analysis was then conducted to evaluate significant differences among species of bee, wasp, and parasite. A Pearson’s R correlation was used to examine whether the intraspecific variation in emergence day, as determined by the standard error of the mean, increased with increasing mean emergence day across the community. Finally, the intertegular (IT) span (Cane 1987; Greenleaf et al. 2007; Williams et al. 2010) was measured (in mm) using an ocular micrometer attached to a dissecting microscope on a sample of 5–10 individuals per species and a linear regression was used to compare IT and mean emergence day among all species. All statistics were completed using the R statistical program v3.2.2 (R Core Team 2015).

## Results

Of the 47 species examined, there were 22 species and six genera of bee in two families (all in the superfamily Apoidea). These included bees in the genus, *Hylaeus* (Colletidae), as well as *Osmia, Heriades, Hoplitis, Chelostoma, Megachile* (all Megachilidae). There were 16 species of cavity-nesting wasp in nine genera and four families, these included *Isodontia* (Sphecidae), *Passaloecus*, *Psenulus* and *Trypoxylon* (Crabronidae), as well as *Ancistrocerus*, *Euodynerus*, *Symmorphus* (Vespidae), and *Auplopus* and *Dipogon* (Pompilidae) (Figure 2). Nine species in seven genera and 5 families of parasites were identified from three orders (Hymenoptera, Coleoptera, Diptera), and parasites, *Ephialtes manifestator* (Linnaeus) and *Anthrax irroratus* Say had more than one host (Table 1).

**Table 1.** Parasite-host associations determined post-emergence as recorded from trap nests in the study region over the three years of sampling. ‘Days’ represents the difference in mean emergence day between the parasite and the host. For parasites having more than one host, ‘Days’ was calculated for each parasite-host pair.

| Parasite | Family | Order | Days | Hosts |
| --- | --- | --- | --- | --- |
| *Sapyga lousi* Krombein | Sapygidae | Hymenoptera | + 7.7 | *Heriades carinata* |
| *Sapyga centrata* Say | Sapygidae | Hymenoptera | + 0.1 | *Osmia pumila* |
| *Ephialtes manifestator* (Linnaeus) | Ichneumonidae | Hymenoptera | - 2.7 | *Passaloecus gracilis*, |
|  |  |  | + 3.1 | *Passaloecus cuspidatus*, |
|  |  |  | - 1.7 | *Trypoxylon frigidum* |
| *Perithous divinator* (Rossi) | Ichneumonidae | Hymenoptera | - 3.6 | *Psenulus pallipes* |
| *Chrysis cembricola* Krombein | Chrysididae | Hymenoptera | - 0.6 | *Symmorphus canadensis* |
| *Caenochrysis doriae* (Gribodo) | Chrysididae | Hymenoptera | + 2.8 | *Trypoxylon frigidum* |
| *Caenochrysis tridens* (Lepeletier) | Chrysididae | Hymenoptera | - 7.7 | *Trypoxylon collinum* |
| *Nemognatha piazzata* (Fabricius) | Meloidae | Coleoptera | - 4.3 | *Megachile rotundata* |
| *Anthrax irroratus* Say | Bombyliidae | Diptera | + 24.2 | *Osmia caerulescens*, |
|  |  |  | + 24.3 | *Osmia pumila* |

Interspecific mean emergence day (e.g. number of incubation days to adulthood) among all 47 species was significantly different (F_47_=786.29, p<0.001) and ranged from 0 to 38 (Figure 2). Many significant differences were noted. For example, the data confirmed one anticipated pattern, that mason bees (*Osmia* spp.) emerge significantly earlier than all others because they overwinter as adults (Fye 1965; Bosch et al. 2001) (Figure 2). The invasive large resin bee, *Megachile sculpturalis*, was the largest species recorded in this study and emerged significantly later than all other species (average emergence day = 37.4±0.7 days; mean±SE) (Figure 2). The greatest overlap (e.g. average emergence day did not differ significantly) occurred between day 12 and 17, when 15 species (three cavity-nesting bees, ten cavity-nesting wasps, and two parasites) emerged (Figure 2).

Parasites had significantly higher variation in adult emergence than both cavity-nesting bees and wasps (F_2_=4.190, p=0.021). The difference in the mean emergence day between parasite and host varied depending on the parasite species. For example, the parasitic fly, *A. irroratus* emerged on average 25 days after its host, whereas emergence day of three cuckoo wasps were more similar to that of their host: on average, *Chrysis cembricola* Krombein emerged on the same day as *Symmorphus canadensis* (de Saussure), *Caenochrysis doriae* (Gribodo) emerged 2.8 days later than its host *Trypoxylon frigidum* Smith, and *Caenochrysis tridens* (Lepeletier) emerged 7.7 days earlier than its host *Trypoxylon collinum* Smith (Figure 2).

Late emerging species exhibited greater intraspecific variation in emergence day. A Pearson’s R correlation showed there was a significantly positive correlation between mean and variation in emergence time (Figure 3). Finally, the average emergence time was positively correlated with body size when all 47 species were included in a linear regression analysis (Figure 4).

**Figure 3.**
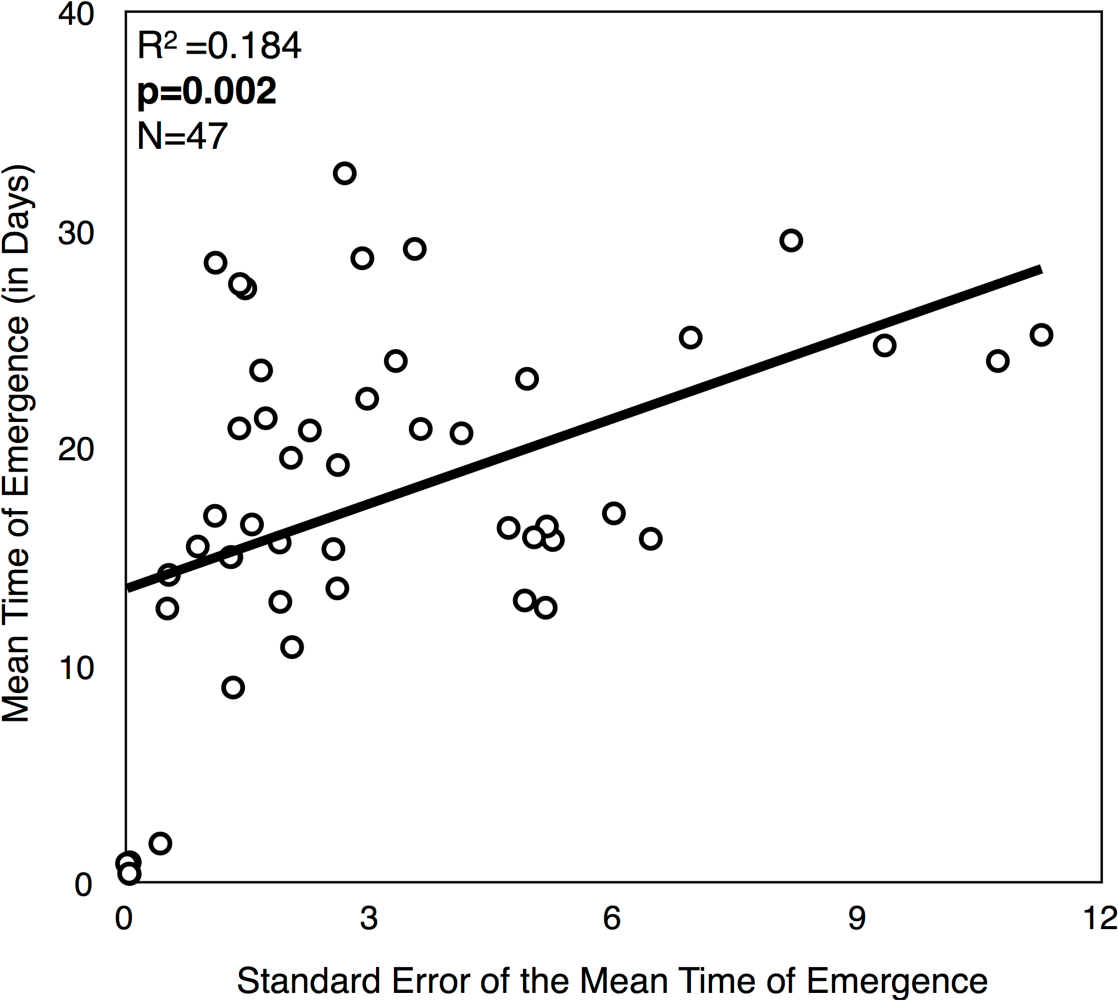
Scatterplot illustrating a significantly positive community-wide correlation between intraspecific mean and variation in emergence time.

**Figure 4.**
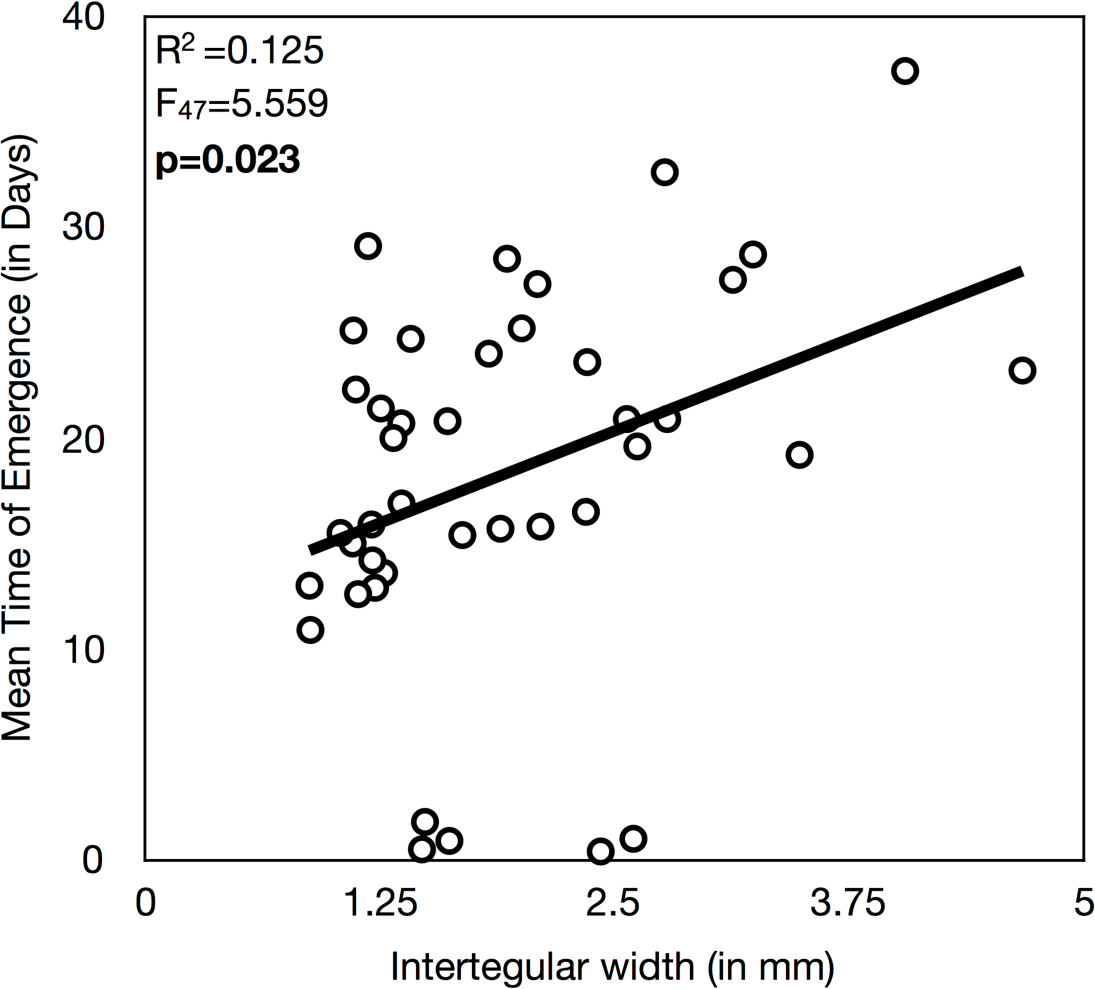
Scatterplot showing the significant relationship between mean emergence time and body size as measured by the intertegular width.

## Discussion

There was a significant interspecific difference in average emergence day among the 47 species of cavity-nesting bees and wasps, and their parasites evaluated. There was evidence to support both hypotheses, that 1) there would be greater intraspecific variation in emergence day in late emerging species compared to early emerging species, and 2) late emerging species would be the largest in body size. Interpreting overlap in emergence time can inform basic knowledge of their species in communities, but also indirectly, potential competition for nesting resources at trap nests. These findings can also support action to enhance target species that use trap nests, by, for example, precisely-timed placement of trap nests in landscapes to increase populations of some species over others.

Bees in the genus *Osmia* and one specialist parasite of *O. pumila* Cresson, the wasp *Sapyga centrata*, emerged significantly earlier than all other species (Figure 2). They become active at the start of spring and this life-history strategy greatly narrow the variation in adult emergence (Figure 3). The remaining bees, wasps, and parasite species use warming temperatures as a cue to initiate transition from pupa to adult, a process which is subject to variation in time to completion due to environmental changes (Tepedino and Parker, 1986; Kemp and Bosch, 2014). Between day 12 and 17 there was an overlap in the mean emergence day among ten cavity-nesting wasp species (Figure 2). These wasp species are each predators of a different group of invertebrates, for example, spiders (*Trypoxylon*; Medler 1967), caterpillars and beetle larvae (*Symmorphus*; Cowan 1981), or aphids (*Passoloecus*; Fricke 1993). Since these wasps were similarly sized, competition for nesting locations in this community could be a more limiting resource than prey opportunities (Wcislo 1996; Potts et al. 2005).

Interspecific body size was positively correlated with mean emergence time and this agrees with other comparative studies that show larger insects take longer to develop (Kingsolver and Huey, 2008). Cavity-nesting bees and wasps select nesting locations with inner widths that are closest to their own body width, and so recording interspecific intertegular widths within a community (see Figure 2) can help practitioners decide on nesting tube dimensions when implementing trap nests to target specific species of interest and known body sizes (Lee-Mäder et al. 2010). Late emerging species exhibited greater intraspecific variation than early emerging species (Figure 3). Being larger can confer a number of benefits; for example, larger bodied solitary bee species can carry more pollen (e.g. Kendall and Soloman 1973) and larger wasps can carry larger prey (e.g. O’Neill 1985; Coelho 1997). However, longer periods of time as prepupae in a nest could increase mortality by parasitism or predation (Stearns and Koella 1986; Blanckenhorn 2000).

The tropical-in-origin and invasive bee, *Megachile sculpturalis*, was the largest species recorded, and the latest to emerge from the nest (Mangum and Sumner 2003). This bee is known to attack and replace a native bee, *Xylocopa virginica* (Linnaeus) (Roulston and Malfi 2012), and likely others coexisting at trap nests. Since *M. sculpturalis* uses tree resins similar to one desirable native bee, *M. campanulae* (Robertson) which emerges significantly earlier (day=27.2±3.3), *M. sculpturalis* can be quickly identified and removed due to this 10+ day difference in emergence time. Knowledge of emergence timing within a community of trap nesting bees and wasps can therefore finetune temporal applications of strategies for supporting target species such as monitoring and removal of invasive species.

Parasite-host association and diversity can inform understanding of parasites as indicators of habitat quality and community-level change (Sheffield et al. 2013). Some authors have noted that parasites are synchronized with hosts and emerge slightly later relative to them (Thorp et al. 1983; Baker et al. 1985). These parasites are typically those that attack larva and so emerging after the host ensures host prey is available. For example, in this study the parasitoid *Anthrax irroratus* attacked two *Osmia* species and emerged 25 days after both hosts, at which point he *Osmia* females had mated and begun to build nests containing the next generation (Table 1). Scott and Strickler (1992) also recorded *A. irroratus* emerging one month after its hosts, *Megachile relativa* Cresson and *M. inermis* Provancher. The parasite larvae develop on the prepupae of the bee host after hatching from a tiny egg ‘flicked’ indiscriminately into the nest by the fly as she hovers in front of the nest entrance (Minckley 1989). Each fly larvae overwinter and develop to adults in the spring (Gerling and Hermann 1976). Cleptoparasites on the other hand replace the host egg with their own, or their early instar larvae kill the host egg or larva, and so have an emergence time that is more similar to that of the host (Forrest and Thomson 2011). In this study, cleptoparasites included *Chrysis cembricola*, *Caenochrysis doriae* and *Caenochrysis tridens,* which all emerged within a week of their host (Figure 2).

Documenting the identity and adult emergence order of coexisting solitary bees, wasps, and parasites in communities can provide significant information about competition and niche overlap in these important and ecologically similar taxa (Frankie et al. 1998; Tscharntke et al. 1998; Bosch and Kemp 2002; Tylianakis et al. 2007; Forrest and Thomson 2011). One limitation of this study is that the emergence times were recorded based on controlled post-incubation temperature and humidity. More work is needed to examine adult emergence of feral populations of solitary bees and compare patterns to those obtained from controlled settings (O’Neill et al. 2010).

Solitary cavity-nesting bees and wasps, that i) compete for a common nesting resource, ii) readily use artificial trap nests, and iii) easily managed by practitioners and citizens intent on enhancing their populations and services they provide, are excellent model organisms for community ecology research, conservation, and outreach with the public on their importance (Lee-Mäder et al. 2010; Colla and MacIvor 2017). These data on interspecific overlap in nesting resources can improve initiatives for pollination service management, and a growing number that are interested in enhancing pest-controlling wasps. Interpreting adult emergence order and overlap in trap nest communities can also support conservation, for example, by knowing when to replace nest tubes with fresh empty ones to ensure adequate supply over the season (MacIvor 2017), or for monitoring invasive species requiring control (Barthell et al. 1998). Trap nests provide a wealth of information on these communities, and so should be in the toolbox of conservation scientists and practitioners working with these critically important insects.

## Acknowledgements

Thanks to Baharak Salehi and Jen Albert for help with the incubation and rearing, also Dr. Laurence Packer and Charlotte de Keyzer for useful comments on the manuscript. Funding was provided by an NSERC-CGS (CGS D 408565) awarded to the first author, and an NSERC discovery grant awarded to the supervisor Dr. Laurence Packer.

